# Types of cis- and trans-gene regulation of expression quantitative trait loci across human tissues

**DOI:** 10.1101/2022.01.24.477617

**Authors:** Jarred Kvamme, MD Bahadur Badsha, Evan A. Martin, Jiayu Wu, Xiaoyue Wang, Audrey Qiuyan Fu

## Abstract

Expression quantitative trait loci (eQTLs) have been identified for most genes in the human genome across tissues and cell types. While most of the eQTLs are near the associated genes, some can be far away or on different chromosomes, with the regulatory mechanisms largely unknown. Here, we study cis- and trans-regulation by eQTLs on protein-coding genes and long noncoding RNAs (lncRNAs) across nearly 50 tissues and cell types. Specifically, we constructed trios consisting of an eQTL, its cis-gene and trans-gene and inferred the regulatory relationships with causal network inference. We identify multiple types of regulatory networks for trios: across all the tissues, more than half of the trios are inferred to be conditionally independent, where the two genes are conditionally independent given the genotype of the eQTL (gene 1 ← eQTL → gene 2). Around 1.5% of the trios are inferred to be mediation (eQTL → mediator → target), around 1.3% fully connected among the three nodes, and just a handful v-structures (eQTL → gene 1 ← gene 2). Unexpectedly, across the tissues, on average more than half of the mediation trios have the trans-gene as the mediator. Most of the mediators (cis and trans) are tissue specific, and cis-gene mediators are significantly enriched for protein-coding genes, whereas trans-gene mediators have a similar distribution of protein-coding genes and lncRNAs to the whole genome.

## 1. Introduction

Gene expression is regulated by genetic variants, among many factors. Expression Quantitative Trait Loci (eQTLs) have been identified to be widespread in the genome by several other consortia [Võsa et al., 2021, The GTEx Consortium, 2020b, Bryois et al., 2014, Battle et al., 2014, Lappalainen et al., 2013, Powell et al., 2012]. In particular, the Genotype-Tissue Expression (GTEx) consortium examines the expression profiles and identifies regulatory variants in around 50 tissues and cell types. They have identified at least one cis-eQTL (i.e., the eQTL is near its target gene, which is the cis-gene of this eQTL) for nearly all the protein-coding genes and nearly 70% of the long intergenic noncoding RNA (lincRNA) genes, as well as a small number of interchromosomal trans-eQTLs (i.e., the eQTL is far away from its target gene, which is the trans-gene of the eQTL). On the other hand, trans-eQTLs are enriched in the GWAS (GenomeWide Association Study) Catalog variants [The GTEx Consortium, 2020b], suggesting potentially important roles of trans-eQTLs in complex traits and diseases.

However, the relationship between the cis- and trans-gene of the same eQTL remains unclear. A popular model is cis-gene mediation, where eQTL regulates the cis-gene (e.g., a transcription factor), which acts as the mediator and regulates the trans-gene [Bryois et al., 2014, Pierce et al., 2014, Kirsten et al., 2015, Yao et al., 2017, Yang et al., 2017, Delaneau et al., 2019, The GTEx Consortium, 2020b, Yang et al., 2021, Viñuela et al., 2021], possibly through regulatory elements (such as cis-regulatory domains, which may contain transcription factor binding sites) on the genome [Delaneau et al., 2019]. However, other modes of relationships have also been observed: Bryois et al. [2014] analyzed 869 lymphoblastoid cell lines with a highly conservative approach and identified 49 cases where the eQTL regulates the cis-gene and the trans-gene independently, and 2 cases where the eQTL affects the trans-gene, which in turn affects the cis-gene, in addition to 19 cis-gene mediation trios. Delaneau et al. [2019] also estimated a high probability of 0.86 for an eQTL to regulate the cis- and trans-gene independently. Furthermore, there is experimental evidence supporting functionally important inter-chromosomal interactions exist. For example, enhancers located on multiple chromosomes may converge and regulate the expression of olfactory receptor genes in cis and in trans [Bashkirova and Lomvardas, 2019, Monahan et al., 2019], suggesting that trans-regulation may be complex and much of it remains unknown.

Analyses of the multiple tissues and cell types of the GTEx consortium have further demonstrated tissue sharing and tissue specificity in the effect of eQTLs: around 80% of all the eQTLs found are shared in more than five tissues, and 25-30% of all the eQTLs are shared in nearly all the GTEx tissues [The GTEx Consortium, 2020b]. To better understand regulation of cis- and trans-genes across tissues, we take a causal network approach to classify possible regulatory relationships in trios of an eQTL and its cis- and trans-target genes, using 48 tissues and cell types in the GTEx version 8 data [The GTEx Consortium, 2020b]. Our analysis goes beyond cis-gene mediation and examines diverse regulatory types, while also accounting for potential confounding from other genes. Our analysis further examines tissue sharing and tissue specificity of these causal relationships. Similar to GTEx, here we focus on protein-coding genes and long noncoding RNAs (lncRNAs) to reduce the impact of potential mappability issues in the RNA-sequencing data [The GTEx Consortium, 2020b].

## 2. Results

### 2.1. Multiple types of regulatory networks are identified for trios

For the causal network analysis here, we used our MRPC method [Badsha and Fu, 2019, Badsha et al., 2021] (version 3.0.0; https://cran.r-project.org/web/packages/MRPC/index.html). MRPC stands for “incorporating the principle of Mendelian Randomization into the PC algorithm", and the PC algorithm is a classical algorithm for inferred directed networks that is named after developers Peter Spirtes and Clark Glymour [Spirtes et al., 2000]). MRPC uses the eQTL as the instrumental variable, and views eQTLs, genes and potential confounding variables as nodes in a network that can represent diverse causal relationships among these nodes.

This approach allowed us to identify multiple types of regulatory networks (or models) for trios (see Section “MRPC analysis accounting for confounding" in Methods). The more interesting types among them are mediation (M_1_), v-structure (M_2_), conditional independence (M_3_, a.k.a., pleiotropy), and fully connected (M_4_) (Figure 1A). In these networks, the two genes have correlated expression levels, although the correlation arises from different regulatory mechanisms. Specifically, in the mediation network, one gene acts as the mediator between the eQTL and the other gene. When conditioned on the mediator, the downstream target is determined only by the mediator and is independent of the eQTL. In the network of conditional independence, the two genes are regulated by the same eQTL, inducing correlation between the two genes, even though there is no direct relationship between them. In the fully connect network, the two genes are not only regulated by the same eQTL, but also influence by additional, unknown processes between the two genes that lead to extra correlation between them. By contrast, in the v-structure network, both the eQTL and a gene are the parents of the other gene. Whereas the eQTL and the “parent" gene are independent, their information becomes dependent when the expression of the “child" gene is given. Apart from these networks, there are also inferred networks with only one edge – these are under the null model (M_0_) category or the “other" category.

**Figure 1.**
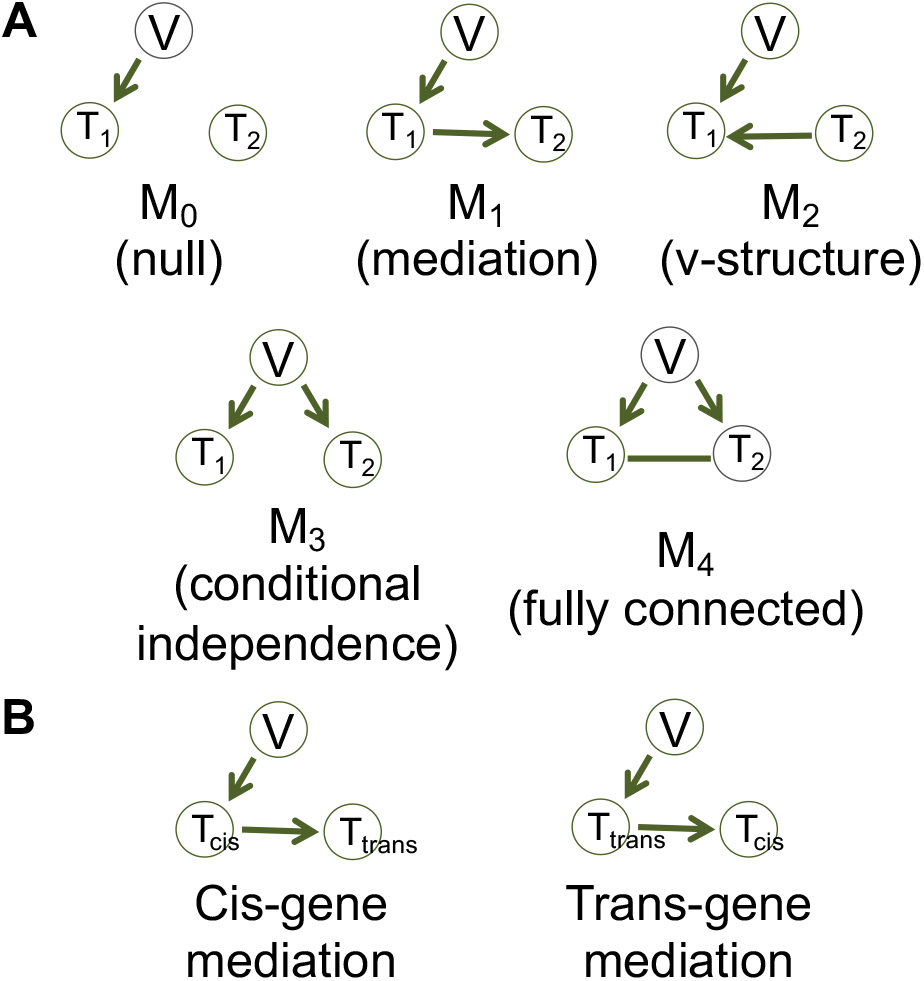
The regulatory relationships of a trio that may be inferred by MRPC. (A) M_0_ through M_4_ are the five basic causal networks that may be detected by our MRPC method. (B) The two types of the mediation model (M_1_): cis-gene mediation and trans-gene mediation.

Across tissues, the number of trios we selected for network inference varies from 627 to 4,668 (median: 2,278; mean: 2,259; Supplementary Table 1). The same eQTLs or genes may appear in multiple tissues. The number of trios generally increases with the sample size (median: 235; mean: 315.2; Supplementary Figure 1). We further derived principal components (PrCs) from the wholegenome gene expression data in each tissue. Using Holm’s method to control the familywise error rate across all the p-values at 5% [Holm, 1979], we identified PrCs that are highly significantly associated with the eQTLs or genes in all the trios of that tissue. We then applied MRPC to each trio, together with the associated PrCs. The number of PrCs included in the trio analysis varies from 0 to 13 across tissues (median: 0-2; Supplementary Table 2).

We applied two different methods, namely the LOND (Levels based on Number of Discoveries) method and the ADDIS (ADaptive DIScarding) method, to control the false discovery rate (FDR) in MRPC (see Section “MRPC analysis accounting for confounding" in Methods). MRPC-LOND tends to be more conservative than MRPC-ADDIS, although they produced qualitatively consistent results here. Below, we present mostly the results from MPRC-LOND in the main text, and include the results MPRC-ADDIS in the supplementary figures and tables.

Among these trios (Figure 2, Supplementary Table 1), 57% are inferred to be conditionally independent (median: 56.2%; mean: 54.9% across tissues), 1.6% to be mediation (median: 1.6%; mean: 1.8% across tissues), 1.3% to be fully connected (median: 1.3%; mean: 1.3% across tissues), and 0.1% to be v-structures (median: 0.1%; mean: 0.1% across tissues). The rest are inferred to have one or no edges. MRPC-ADDIS inferred a similar percentage for conditionally independent trios (58%) and for mediation trios (1.8%), but more fully connected trios (5.2%) and v-structures (0.6%) (Supplementary Figure 2, Supplementary Table 3).

**Figure 2.**
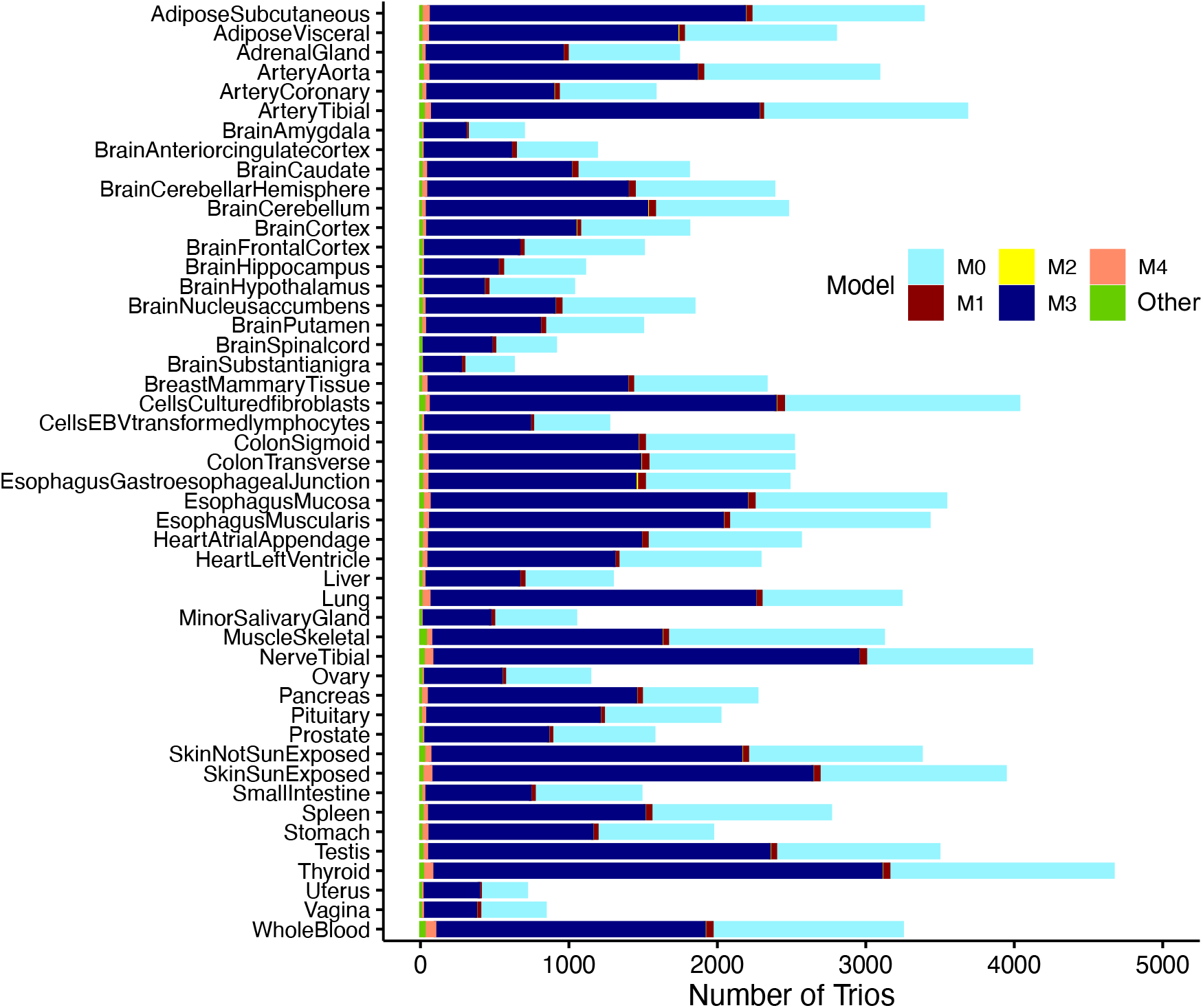
A stacked bar plot of inferred regulatory types of trios across GTEx tissues and cell types. Each causal network (or model) is represented by a unique color. “Other" refers to inferred networks that are not any of the five basic models.

### 2.2. Cis- versus trans-gene mediation

The mediation model is of particular interest, as in general the hypothesis is that an eQTL regulates its trans-gene target through the cis-gene, which acts as a mediator. We term this the “cis-gene mediation" (Figure 1B). A trans-gene may also act as a mediator; we term this type of mediation the “trans-gene mediation" (Figure 1B).

Unexpectedly, we inferred a large number of trios to be trans-gene mediation (Figure 3; Supplementary Tables 4, 5, 6, 7 and 8). Among the 1,718 mediation trios we inferred across tissues, we identified 395 (23%) cis-gene mediation trios and 1,323 (77%) trans-gene mediation trios. The number of cis-gene mediation trios varies from 3 to 15 across tissues (median: 8; mean: 8.2), and that of trans-gene mediation trios varies from 7 to 46 (median: 28; mean: 27.6). Furthermore, the 395 cis-gene mediation trios were also inferred by MRPC-ADDIS (with less stringent FDR control). Only 696 of the 1,323 trans-gene mediation trios were also inferred by MRPC-ADDIS; most of the remaining trios (491) were inferred to be M_4_, which contains three edges.

**Figure 3.**
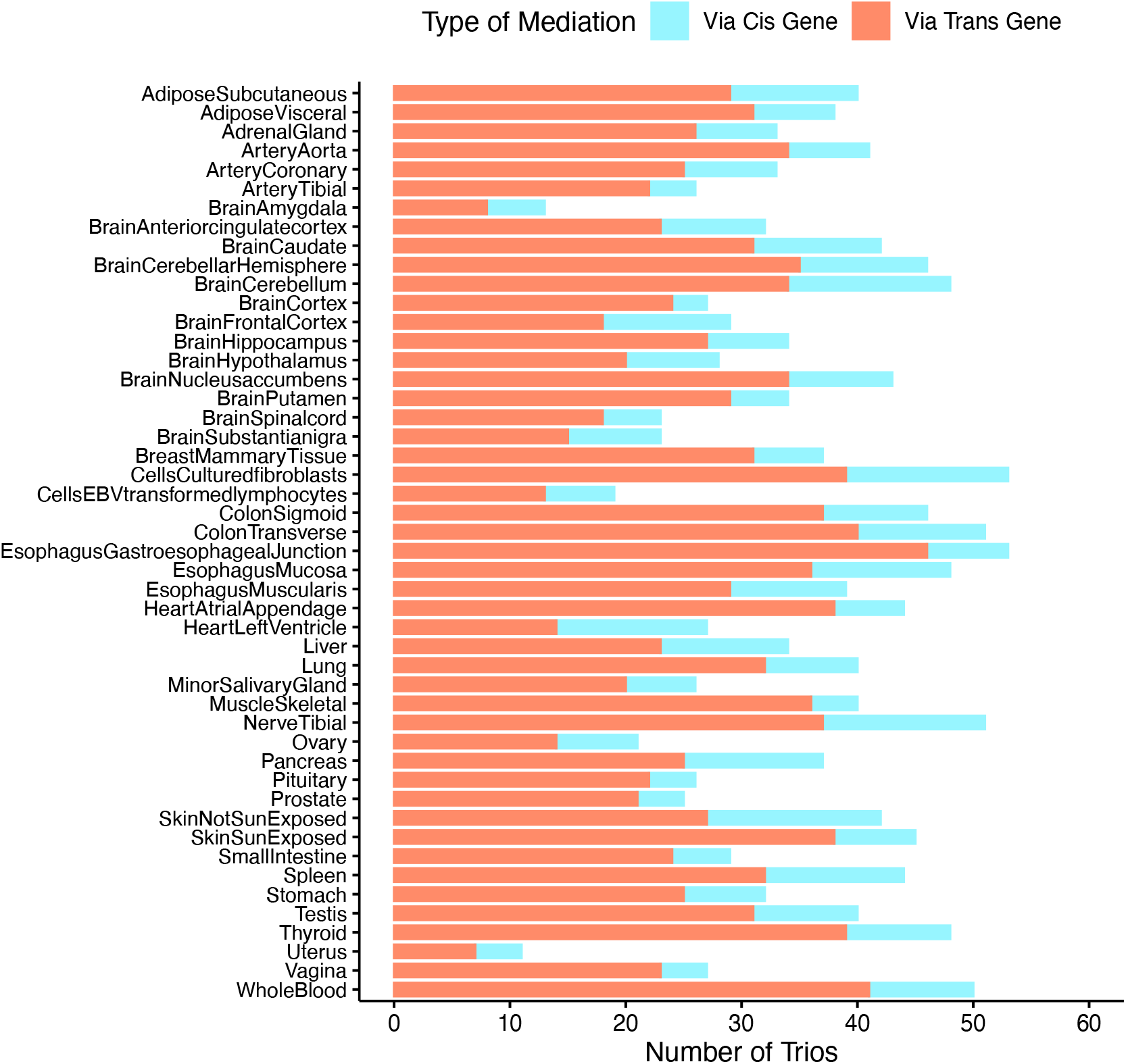
A stacked bar plot of inferred mediation types across GTEx tissues and cell types.

In cis-gene mediation trios, the absolute Pearson correlation between an eQTL and its cis-gene tends to be larger than that between the eQTL and its trans-gene (Figure 4). This trend is reversed in trans-gene mediation trios, which is consistent with the statistical inference: direct association (i.e., having an edge between two nodes) tends to have a stronger correlation than indirect association (i.e., not having an edge between two nodes), and an edge with a stronger association is more likely to be causal than an edge with a weaker association. Due to the same reasons, when the cis-gene is inferred to be the mediator, the association between the two genes also tends to be stronger than that between the eQTL and the trans-gene. When the trans-gene is inferred to be the mediator, the association between the two genes tends to be stronger than that between the eQTL and the cis-gene.

**Figure 4.**
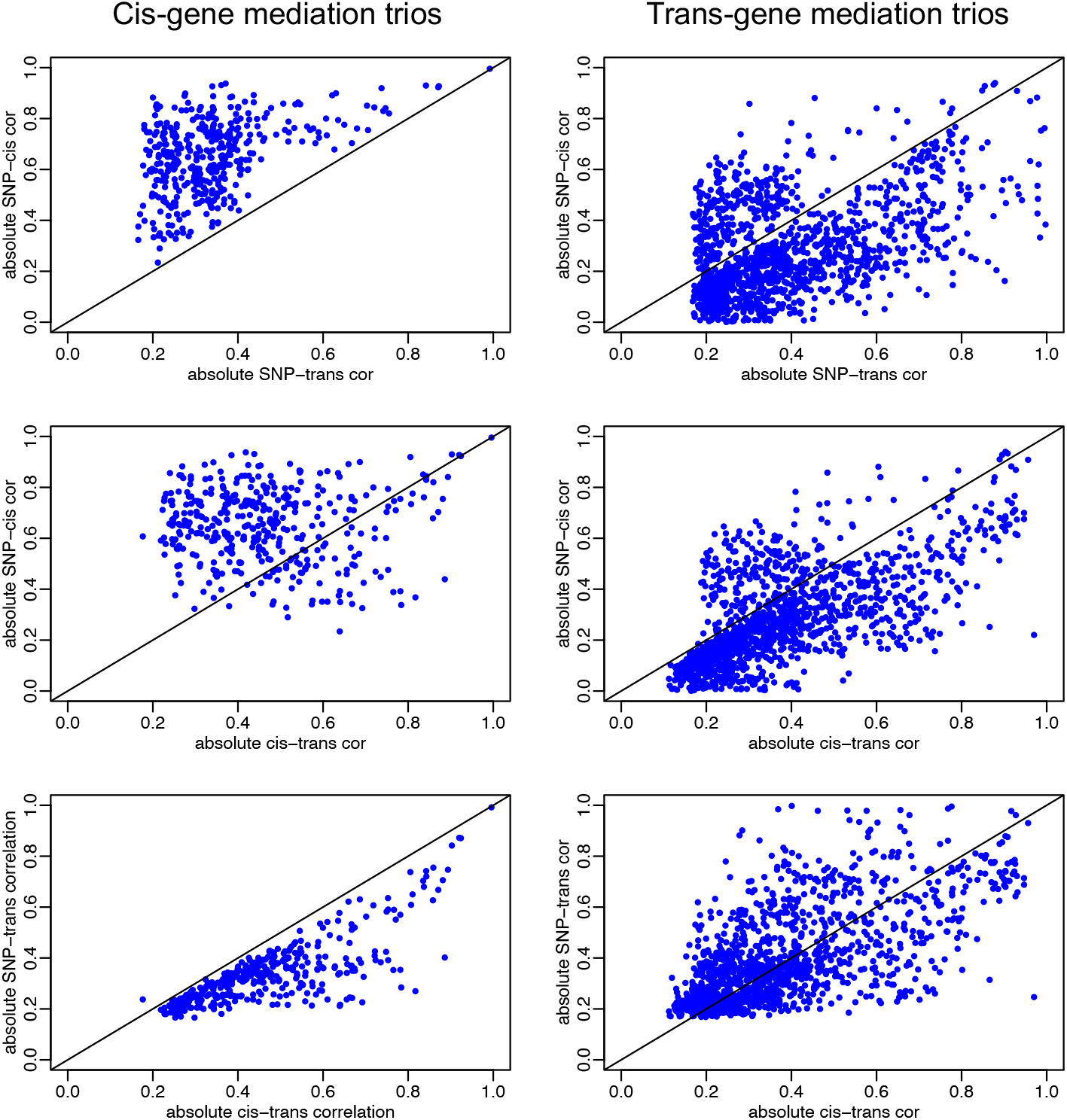
Pearson correlations in mediation trios. Top: scatterplots of the absolute correlations between the SNP genotype and the cis-gene expression versus that between the SNP and the trans-gene expression. Middle: scatterplots of the absolute correlations between the SNP and the cis-gene expression versus that between the expression of the two genes. Bottom: scatterplots of the absolute correlations between the SNP and the trans-gene expression versus that between the expression of the two genes.

These mediation trios involve 1,339 unique genes as mediators. Among them, 243 are cis-genes and not trans-genes for an eQTL, 1,066 are trans-genes and not cis-genes, and 30 are both cis- and trans-genes for an eQTL (Supplementary Table 9). Most of these mediators appear to be tissue specific: 215 (79%) of the 243+30=273 cis-gene mediators appear only in one tissue, whereas 1,041 (95%) of the 1,066+30=1,096 trans-gene mediators appear only in one tissue. (Figure 5). The results from MPRC-ADDIS also show that most mediators are tissue specific (Supplementary Figure 5).

**Figure 5.**
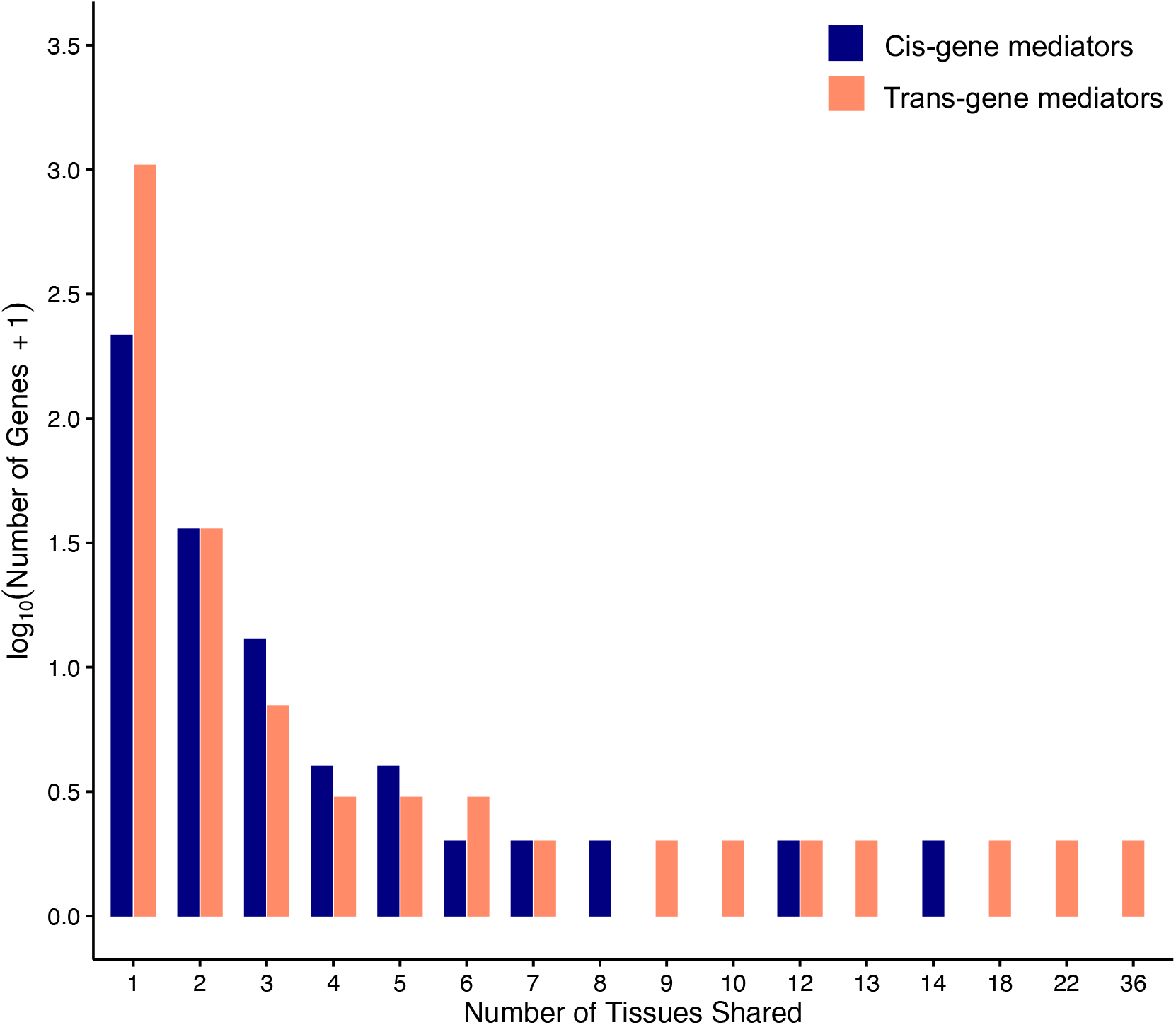
Histograms of tissue sharing for cis-gene and trans-gene mediators.

We next investigated whether the mediators are enriched for protein-coding genes or lncRNAs, using the Ensembl human gene annotations (Figure 6; Supplementary Table 9). Among 273 cis-gene mediators, 201 (74%) are protein-coding genes and 72 (26%) lncRNAs. Among 1,096 trans-gene mediators, 618 (56%) are protein-coding genes and 478 (44%) lncRNAs. Of the 43,778 protein-coding genes and lncRNAs in the human genome, protein-coding genes account for 57% and lncRNAs 43%. The distribution of the two types does not differ between trans-gene mediators and the whole genome. On the other hand, cis-gene mediators are significantly enriched for protein-coding genes, compared to trans-gene mediators (chi-square test p-value: 5×10^−7^) and to the whole genome (chi-square test p-value: 2×10^−7^). The mediators inferred by MRPC-ADDIS show a similar distribution of protein-coding genes and lncRNA, and the corresponding chi-square test p-values are similarly small (1×10^−10^ and 2×10^−11^, respectively).

**Figure 6.**
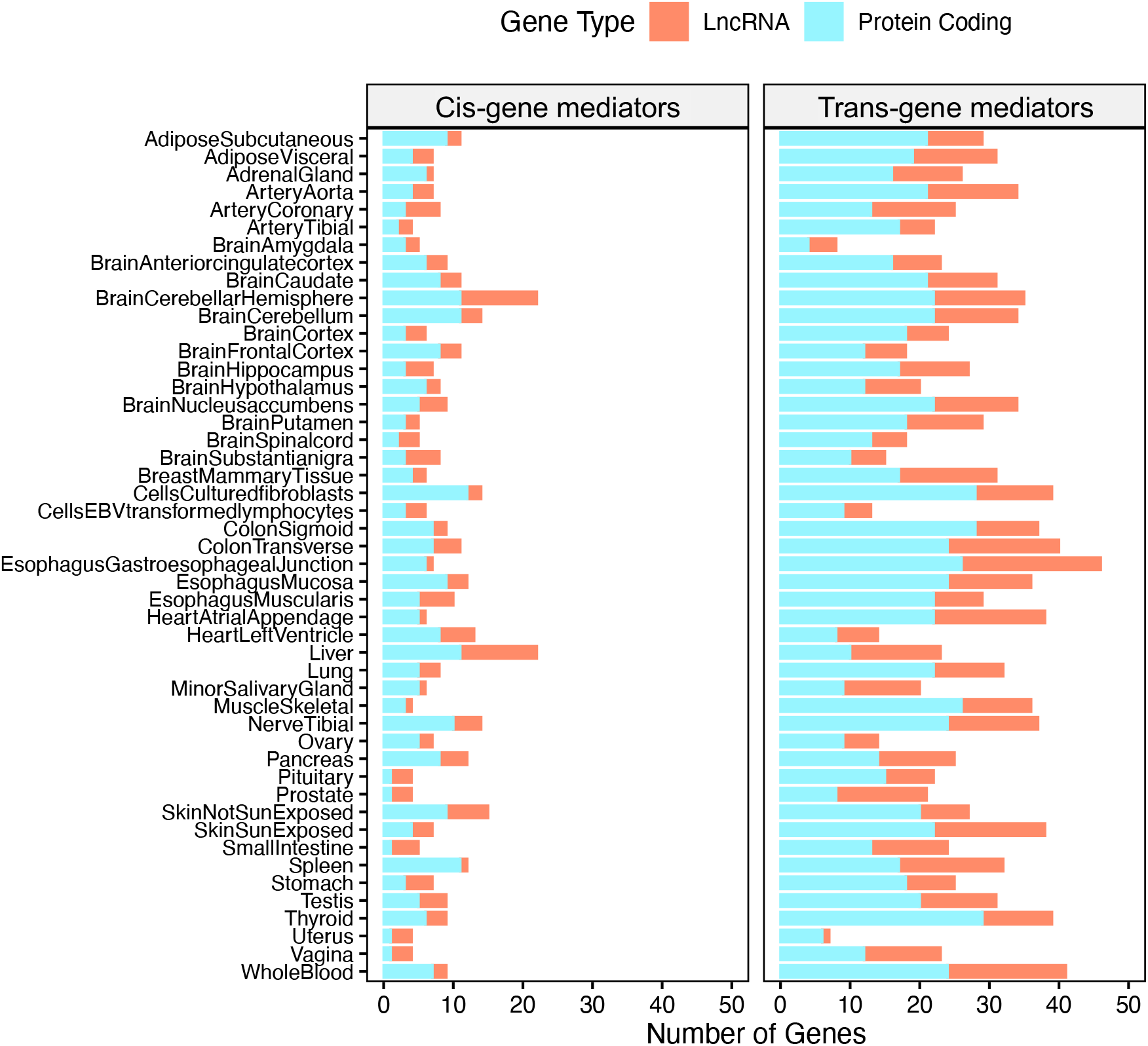
A stacked bar plot of mediators by gene types across GTEx tissues and cell types.

### 2.3. Analysis of HiC sequencing data for trios of trans-gene mediation

We examined the HiC sequencing data for physical interactions in multiple cell types (lung, skin, lymphoblastoid cells, and fibroblast cells; Rao et al. [2014], Sanborn et al. [2015], Nir et al. [2018]; see “HiC sequencing data analysis" in Methods) for potential evidence that may support trans-gene mediation. HiC sequencing reads indicate that two loci on the genome are physically close in space, even when the loci are far apart on the linear genome [Lieberman-Aiden et al., 2009]. Studies of promoter-enhancer interactions have suggested that enhancers that may be located far away from a gene can be brought close to the promoter of this gene through chromatin looping, thus facilitating the regulation of gene expression [Bashkirova and Lomvardas, 2019, Monahan et al., 2019]. Here, we were interested in regulation of an eQTL and its trans-gene mediators. The hypothesis is that trans-regulation may involve physical contact between the eQTL and the gene, which may be captured by HiC sequencing. However, we did not observe significant enrichment of HiC reads for any trio of trans-gene mediation (Supplementary Tables 10 and 11).

For example, in fibroblasts (Supplementary Table 10), we were able to extract as many as 4,287 HiC reads between a 200kb neighborhood of the eQTL at chr14:97,718,183 and a 200kb neighborhood of its trans-gene *BCL11B* (*BCL11* transcription factor B) also on chr14 at chr14:99,169,287-99,272,197 (1.45 Mb away from the eQTL). When we performed Monte Carlo simulation to generate the null distribution of the HiC reads for chromosome 14, we obtained a Monte Carlo p-value of 0.01 (with a q-value of 0.10). In skin unexposed to the Sun (Supplementary Table 10), between the eQTL at chr17:28,499,368 and its trans-gene *TSPO* at chr22:43,151,547-43,163,242, we were able to extract only 2 reads, although low numbers of interaction reads are common between two chromosomes. The Monte Carlo p-value was 0.27 with a q-value of 1.00.

### 2.4. Comparing MRPC to GMAC on the mediation trios

To further validate the observation of many trans-gene mediation trios, we applied the GMAC (Genomic Mediation analysis with Adaptive Confounding adjustment) method [Yang et al., 2017] to all the candidate trios in five GTEx tissues with the largest sample sizes: adipose subcutaneous, tibial artery, muscle skeletal, sun exposed skin, and whole blood. GMAC aims to detect mediation; specifically, GMAC examines in each trio whether the data supports the candidate mediator having an effect on the target gene, even if the eQTL may also have a direct effect on the target (see Discussion on the implications of this approach). Although Yang et al. [2017] considered only cis-genes to be candidate mediators in their analysis, the GMAC method itself is agnostic to which gene can be the candidate mediator. In our application, we applied GMAC to each trio twice: first treating the cis-gene as a potential mediator, and next treating the trans-gene as a potential mediator. With this analysis, we were interested in testing whether a method different from MRPC also identified many trans-gene mediation trios.

At an FDR of 10%, GMAC identified on average 137 (median: 124) cis-gene mediation trios and 120 (median: 120) trans-gene mediation trios across the five tissues (Supplementary Table 12). Most of these trios are the same trios, and GMAC inferred mediation when the cis-gene was the mediator and also when the trans-gene was the mediator. This means that these trios follow an M_4_ model in our framework where the edge between the two genes is bidirected (Figure 1). Nevertheless, this result is consistent with ours and confirms that trans-gene mediation is at least as common as cis-gene mediation.

## 3. Discussion

Using our causal network inference method, we identified multiple types of regulatory relationships in trios of an eQTL and its cis- and trans-genes, which provides a more comprehensive picture of the complex relationships in trios. Across the tissues, more than half of the tested trios are inferred to be conditionally independent, and around 1.6% of the trios are inferred to be mediation. Interestingly, on average more than half of the mediation trios have the trans-gene as the mediator. Furthermore, cis-gene mediators are enriched for protein-coding genes compared with the genome average and with the trans-gene mediators.

An edge inferred by our statistical method indicates that the relationship is strong enough not to be explained away by other factors we have considered. Such a strong relationship, albeit still a statistical result, is therefore more likely to reflect a genuine regulatory relationship. On the other hand, it is very likely that an inferred edge condenses a complex regulation process, which may involve many genes or processes other than transcription.

However, although the number of trios we examined is on the order of thousands across tissues, this is still a very small number of possible trios for the human genome. The trios we considered here generally have strong association. This is because when multiple SNPs were identified to be eQTLs for the same cis-gene, we used the SNP with the smallest p-value. As a result, we may have missed many eQTLs that could have slightly weaker association than the chosen one, but may have stronger association with trans-genes. The distribution of the regulation types may be biased due to this omission, but the distribution from our analysis is quantitatively comparable to that from other studies [Bryois et al., 2014, Delaneau et al., 2019], with the majority of the trios being the conditional independence type, and a small percentage of mediation trios.

Several studies have examined cis-gene mediation [Bryois et al., 2014, Yang et al., 2017, The GTEx Consortium, 2020b]. In particular, Yang et al. [2017] systematically identified trios of cis-gene mediation across GTEx tissues (using data from version 7), using their GMAC method, which is also based on the principle of Mendelian randomization and accounts for confounding variables. However, the GMAC method focused exclusively on mediation and cannot detect other relationships. Yang et al. [2017] also focused on trios with strong associations and examined a smaller number of trios in each tissue (median: 1,112.5; mean: 1,473). On the other hand, at an FDR of 5%, they identified more trios of cis-gene mediation than we did in more than half of the tissues: specifically, they identified a median of 102.5 (mean: 140.0) cis-gene mediation trios, much higher than ours. Take whole blood for example, they identified 281 trios of cis-gene mediation, whereas we identified only 50 mediation trios of both types (9 cis-gene mediation trios and 41 trans-gene mediation trios).

Although surprising initially, the observation of a large number of trans-gene mediating trios is consistent with existing literature on the prevalence of trans-regulation. As summarized by Liu et al. [2019], trans-eQTLs contribute to 60-90% of the heritability in gene expression across multiple studies [Price et al., 2008, 2011, Wright et al., 2014, Liu et al., 2017], suggesting that eQTLs often regulate genes that are far away. Trans-gene mediation has also been identified before by other studies. For example, Bryois et al. [2014] applied the CIT (Causal Inference Test) method to the lymphoblastoid cell lines (LCLs) from a cohort of 869 individuals and identified 19 trios of cis-gene mediation, 2 trios of trans-gene mediation, as well as 49 conditional independent trios. The number of trans-gene mediation trios is low in this study, but the number of conditional independent trios is also low, suggesting lower power to detect any type. Overall, the observation that the conditional independent trios are much more frequent than mediation trios is consistent with our findings.

What is the potential mechanism for trans-gene mediation? GTEx found enrichment of trans-eQTLs at *CTCF* (CCCTC-binding factor) binding sites, and hypothesized that such eQTLs may disrupt CTCF binding, which influences the spatial chromatin interaction and therefore gene expression [The GTEx Consortium, 2020b]. Our analysis of the HiC data was inspired by the studies on distal enhancers for olfactory receptor genes [Bashkirova and Lomvardas, 2019, Monahan et al., 2019], and assumed direct contact between an eQTL and its trans-gene. Not observing significant HiC enrichment in our analysis is not necessarily evidence against trans-mediation. The number of reads connecting two chromosomes is generally low, and so is the number of reads connecting two distal regions on the same chromosome. The inconclusive result therefore may have two interpretations: i) There is genuinely no physical interaction; and ii) Physical interaction is not captured by current technology. We currently do not have knowledge of which interpretation is more likely. Furthermore, because of the sparse read counts between distal locations in existing HiC data, we examined a wide neighborhood of each eQTL in our analysis, and this neighborhood of 200kb would generally include the cis-gene of the eQTL. The reads we identified between an eQTL and a trans-gene could also be between the cis-gene and the trans-gene, or between other genes in the two neighborhoods. Võsa et al. [2021] also examined the HiC enrichment between eQTLs (or equivalently cis-genes) and trans-genes, and found significant enrichment compared to all possible gene pairs at the genome level (*p* = 2.4 × 10^−153^), which provides global support for spatial interaction, although it remains difficult to pinpoint which gene pairs are enriched for such interaction.

We validated the common presence of trans-gene mediation with the GMAC method. However, it is important to note that GMAC detects mediation by testing for association between two genes in a trio, in the presence of an eQTL and confounder variables. This means that GMAC tests only the edge between the two genes, regardless the presence of other edges [Yang et al., 2017]. Therefore, mediation under GMAC corresponds to any of three models under our framework: M_1_, M_2_, or M_4_ (all have an edge between two genes), whereas lack of a mediation relationship under GMAC corresponds to either M_0_ or M_3_ (no edge between two genes) (Figure 1). For the trios from the five tissues analyzed by both GMAC and MRPC-LOND, we found that the agreement is as high as 98% between the two methods for no mediation across tissues (Supplementary Table 13). On the other hand, this agreement is only 25-44% for mediation, although this is likely due to MRPC-LOND being a high conservative method (Supplementary Table 14).

The comparison of MRPC and GMAC raises the question: what does mediation mean? GMAC allows an eQTL to regulate a target gene directly *and* through a mediator, whereas MRPC requires that all the eQTL effect goes through the mediator and that there is no edge connecting the eQTL and the target gene. We may consider the former *partial* mediation and the latter *complete* mediation. Partial mediation, however, is a challenge in statistical inference. Consider the following two models: i) *V → T*_1_, *V → T*_2_, *T*_1_ → *T*_2_; and ii) *V → T*_1_, *V → T*_2_, *T*_1_ ← *T*_2_. These two graphical models always have the same likelihood, which means that they are Markov equivalent [Verma and Pearl, 1990] and cannot be distinguished without additional biological information. Both models are interpreted as mediation by GMAC, but are considered to be statistically unidentifiable under MRPC and are therefore inferred as M_4_ (*V → T*_1_, *V → T*_2_, *T*_1_ −*T*_2_; Figure 1) instead.

## 4. Methods

### 4.1. The GTEx genotype and gene expression data used for this study

We used the association test results from GTEx version 8 (in *.egenes.txt.gz, where * refers to a tissue or cell type name) to identify the top eQTL for individual genes [The GTEx Consortium, 2020a]. In the files *.egenes.txt.gz, a single variant is reported for each gene, even though the association may not be strong. We extracted the genes with an association q-value ≤ 0.05, which resulted in 4,934 genes.

### 4.2. PEER normalization of GTEx gene expression data

We used the transcript TPM (transcript per million) data for gene expression in 48 tissues (or cell types). A handful of tissues were removed due to a low sample size (< 100). The sample size of the tissues analyzed here ranges from 114 to 706, with a mean of 315 and a median of 235. Following the standard procedure adopted by the GTEx consortium, we performed PEER normalization (probabilistic estimation of expression residuals) [Stegle et al., 2012] on the whole-genome gene expression data for each tissue. We included the covariates provided by GTEx: sex, platform, PCR, and the top five principal components from the genome-wide genotype data, which may contain signals on the potential population structure. We added age to this list of covariates. Additionally, we included different numbers of PEER factors for each tissue, depending on the sample size: 15 factors for < 150, 30 for 150 − 250, 45 for 250 − 350, and 60 for ≥ 350.

### 4.3. Selection and identification of trios

Using the eQTLs reported by GTEx and the PEER-normalized gene expression described in the two sections above, we next ran the R package MatrixEQTL [Shabalin, 2012] to look for trans-genes located 1 Mb away from the eQTLs with a p-value < 10^−5^. Multiple trans-genes may be identified for the same eQTL. We constructed trios for each eQTL with a cis-gene and a trans-gene. Different trios may have the same eQTL, or the same gene. A gene may also be a cis-gene in one trio but a trans-gene in another. From these trios, we further selected those that have only protein-coding genes or lncRNAs.

### 4.4. Identification of associated confounding variables

We performed principal component analysis on the PEER-normalized, genomewide gene expression in each tissue and then tested the significance of the three variables in each candidate trio to each PrC using a simple regression and obtained a p-value. We used Holm’s method to control the familywise error rate across all the p-values at 5% [Holm, 1979]. Each PrC reflects potential impact from a large number of genes and represents the influence from the larger gene regulatory network to which a trio may belong. We then included these PrCs as additional nodes in the MRPC analysis of the trio. Due to the strong control of Holm’s method, the median number of PrCs included in the end varies between 0 and 2 across tissues.

### 4.5. MRPC analysis accounting for confounding

We used our R package MRPC [Badsha and Fu, 2019, Badsha et al., 2021] to perform the network inference on the individual-level data for each trio in each tissue, including the associated principal components as additional nodes. MRPC builds on the classical PC algorithm (named after its developers Peter Spirtes and Clark Glymour; Spirtes et al. [2000]) for inference of directed acyclic graphs, and incorporates the principle of Mendelian randomization (PMR) [Davey Smith and Hemani, 2014]. As discussed in our earlier work [Badsha and Fu, 2019, Badsha et al., 2021], one class is the general-purpose network inference methods, such as the PC algorithm and its variants (e.g., methods implemented in R packages bnlearn [Scutari, 2010] and pcalg [Kalisch et al., 2012]). Existing methods in this class are computationally efficient but difficult to modify to account for the PMR. The other class is developed for genomic data and explicitly accounts for the PMR (e.g., CIT from Millstein et al. [2009], QPSO from Wang and van Eeuwijk [2014], and findr from Wang and Michoel [2017]). However, the types of causal networks detected by existing methods are often limited to a subset of the five basic models in Figure 1. Recently, several methods [Howey et al., 2020, Yazdani et al., 2016, Zhu et al., 2012], including our MRPC method, have been developed to combine the strengths of two classes of methods for causal network inference.

The PC algorithm consists of two main steps: inferring a graph skeleton, where the key edges are retained but undirected; and determining the direction of the edges. Our MRPC improves both steps and achieves better power and lower FDR on small networks [Badsha and Fu, 2019, Badsha et al., 2021]. The improvement in Step 1 is further explained below. Step 2 in MRPC incorporates the PMR, which takes advantage of the additional information in eQTLs. Under the PMR, the genotypes can be reasonably assumed to be randomly allocated in a natural population, and can therefore be viewed as randomization of the individuals. Since the genotypes influences the phenotypes, but not the other way around, the PMR then views an eQTL as an instrumental variable for causal inference. Causal inference on a trio aims to infer a three-node network for the eQTL and the two genes, with an edge pointing from the eQTL to one or both genes (Figure 1).

Step 1 in MRPC, as well as other PC-like algorithms, starts from a fully connected network and performs a series of tests on each edge to see whether the two nodes are marginally correlated or conditionally correlated, given one other node, or two other nodes, or any subset of other nodes. If a test produces an insignificant p-value, it means that the correlation between the two nodes may be not strong enough or can be explained away by other nodes. The edge would be removed and never tested again. Hypothesis testing in PC-like algorithms is therefore online, meaning that the number of statistical tests to be performed is unknown in advance, and that the threshold for a p-value to be considered significant cannot be fixed beforehand. Several methods have been developed to control the overall FDR in this online setting. We have implemented such a method called LOND [Javanmard and Montanari, 2015] in MRPC, and demonstrated that LOND achieved better power and lower actual FDR than existing methods on small networks through extensive simulations [Badsha and Fu, 2019, Badsha et al., 2021].

However, LOND may be too conservative and leads to the true edges being missed, especially in larger networks [Badsha et al., 2021]. We have therefore also implemented the ADDIS method [Tian and Ramdas, 2019], another less-conservative online FDR control method, in MRPC. We used both ADDIS and LOND in MRPC to control the FDR at 5% for each trio.

To compare the performance of ADDIS and LOND, we performed the following simulation. We included three known confounders in GTEx gene expression data: PCR, platform, and sex. We further used the PrCs derived from the GTEx whole blood gene expression data as additional, potential confounders. We simulated data independently for trios, where each trio has a set of randomly selected confounders. We generated data for a total of 200 trios under all the five basic causal networks (34-47 trios per network) in Figure 1A, with the confounders being the parent nodes of both gene nodes in each trio. We varied the number of confounders, effect size, standard deviation in the error, and minor allele frequency of the eQTL in the simulation. Most of the values used for the parameters are similar to those in the GMAC paper [Yang et al., 2017]. Specifically, the effect size from the eQTL is drawn from Unif (0.5, 1.5), and the effect size from a gene is drawn from Unif (0.5, 1). The standard deviation in the random error is set to be *aβ*, where *β* is the effect size of a gene, and *a* is drawn from Unif (0.3, 1.5). The minor allele frequency is drawn from Unif (0.01, 0.5). For each trio, we selected a random number of PrCs from a discrete uniform distribution between 1 and 15. The effect size from these PrCs was drawn from Unif (0.15, 0.5). The effect size from the three known confounders was drawn from Unif (0.01, 0.1). We then applied MRPC-ADDIS and MRPC-LOND to each dataset. We evaluated recall and precision in two ways: i) of the edges in the true and inferred networks; and ii) only for the edge between the two genes.

This simulation showed that ADDIS and LOND have similar performance overall (Supplementary Table 15). ADDIS achieved better recall (i.e., power) than LOND did across models, although the recall rates were not high. Both ADDIS and LOND achieved similarly high precision rates across models, once again confirming our previous observations that MRPC is conservative but accurate on inferred edges [Badsha and Fu, 2019, Badsha et al., 2021].

### 4.6. Gene type enrichment analysis among mediators

Both cis-genes and trans-genes may be inferred to be the mediator. We used the Ensembl human gene annotations (GRCh38/hg38) to label each gene as a protein-coding gene or a lncRNA. We summarized the counts for cis-gene mediators, and separately for trans-gene mediators. We performed chi-square tests to compare the distribution of gene types among cis-gene (or trans-gene) mediators to the whole genome.

### 4.7. HiC sequencing data analysis

We downloaded four HiC-sequencing datasets from the ENCODE consortium: lung (ENCFF366ERB; Rao et al. [2014], Sanborn et al. [2015]), skin (ENCFF569RJM; Rao et al. [2014], Sanborn et al. [2015]), lymphoblastoid cells (ENCFF355OWW; Rao et al. [2014], Sanborn et al. [2015]), and fibroblast cells (ENCFF768UBD; Nir et al. [2018]). We again used the Ensembl human gene annotations (GRCh38/hg38) to determine the positions of genes. We used the package *StrawR* [Durand et al., 2016] to extract reads from positions along the chromosomes corresponding to the SNP and trans-mediated gene. For all extractions, the resolution, defined as the bin size in the package, was set to 10 kb, which was the finest resolution shared by all four tissues.

We calculated a Monte Carlo p-value to identify whether an observed number of interactions between a SNP and a trans-gene mediator was significant. These p-values were formulated from the upper tail probability of the observed number of reads relative to the empirically generated null distribution. To construct the empirical null distribution for each pair, we randomly drew 10,000 pairs of neighborhoods of 200 kb uniformly located on both the chromosome of the SNP and the chromosome of the trans-gene. The numbers of interaction reads in these neighborhood pairs then constitute the null distribution. The Monte Carlo p-value is then the proportion of reads exceeding the observed number of interaction reads. To account for multiple testing, we applied Holm’s method [Holm, 1979] to control the family-wise error rate at 5%, and the Benjamini and Yekutieli method [Benjamini and Yekutieli, 2001] and the q-value method [Storey, 2002] to control the FDR at 5%.

### 4.8. Comparison with GMAC

To compare the GMAC and MRPC methods, we applied GMAC (version 3.0; https://cran.r-project.org/web/packages/GMAC/index.html) to each of the top five GTEx tissues by sample size. Note that GMAC can detect only mediation but not other relationships. Following the instructions in the GMAC package and consistent with the application in Yang et al. [2017], we used all the principle components of the genomewide expression matrix as the covariate pool, and three additional known covariates in GTEx: the PCR used, the platform used, and sex of the individual in each sample.

For each trio, GMAC identified and removed covariates that were a common child or intermediate variable to the two genes at an FDR of 10%, and identified confounders (defined as a parent node to the two genes in GMAC) at an FDR of 5%. The input to GMAC consisted of the eQTL and the PEER-normalized expression values of the cis- and trans-gene. Since the genotypes are missing in some individuals, we performed imputation using multiple correspondence analysis (MCA; Josse et al. [2016]) prior to the GMAC analysis. We ran GMAC twice on each trio, first with the cis-gene as the potential mediator and second with the trans-gene as the potential mediator. GMAC output a p-value for each trio. Yang et al. [2017] used these unadjusted p-values in two ways to select mediation trios: i) applying a cutoff *p* < 0.05, which means no correction for multiple testing; and ii) applying the q-value method and setting a cutoff of *q* < 0.05 [Yang et al., 2017]. Here, we took the middle road: we applied the q-value method but set the cutoff *q* < 0.1 for each tissue.

We further applied GMAC to the simulated data generated in Section “MRPC analysis accounting for confounding" to compare with MRPC. Since GMAC can infer only M_1_, we assessed recall and precision only on the edge (or its absence) between the two genes (Supplementary Table 16). GMAC showed a very high recall rate (0.97) but low precision (0.79) for this edge, indicating its tendency of inferring the presence of such an edge, regardless whether other edges are present or not. In contrast, MRPC-ADDIS and MRPC-LOND both had low recall (0.42 and 0.33, respectively) but high precision (0.85 and 0.87, respectively), again confirming that MRPC is much more conservative in its inference but tends to get it right.

## Data access

Data from The Genotype-Tissue Expression (GTEx) Project version 8 were obtained from db-GaP (http://www.ncbi.nlm.nih.gov/gap) through accession number phs000424.v8.pht002741.v8.p2. Open-source R packages used in this manuscript include MRPC (version 3.0.0; https://cran.r-project.org/web/packages/MRPC/index.html) and GMAC (version 3.0; https://cran.r-project.org/web/packages/GMAC/index.html). Detailed results generated by our analyses are included in the supplementary material.

## Competing interest statement

The authors declare no competing interests.

## Acknowledgements

We acknowledge the extensive computing support provided by the Research Computing and Data Services from the Institute for Interdisciplinary Data Sciences at the University of Idaho.

This research is supported by the National Institutes of Health (NIGMS P20GM104420).

The Genotype-Tissue Expression (GTEx) Project was supported by the Common Fund of the Office of the Director of the National Institutes of Health. Additional funds were provided by the NCI, NHGRI, NHLBI, NIDA, NIMH, and NINDS. Donors were enrolled at Biospecimen Source Sites funded by NCI/Leidos Biomedical Research, Inc. subcontracts to the National Disease Research Interchange (10XS170), Roswell Park Cancer Institute (10XS171), and Science Care, Inc. (X10S172). The Laboratory, Data Analysis, and Coordinating Center (LDACC) was funded through a contract (HHSN268201000029C) to the The Broad Institute, Inc. Biorepository operations were funded through a Leidos Biomedical Research, Inc. subcontract to Van Andel Research Institute (10ST1035). Additional data repository and project management were provided by Leidos Biomedical Research, Inc.(HHSN261200800001E). The Brain Bank was supported supplements to University of Miami grant DA006227. Statistical Methods development grants were made to the University of Geneva (MH090941 & MH101814), the University of Chicago (MH090951,MH090937, MH101825, & MH101820), the University of North Carolina - Chapel Hill (MH090936), North Carolina State University (MH101819), Harvard University (MH090948), Stanford University (MH101782), Washington University (MH101810), and to the University of Pennsylvania (MH101822). The datasets used for the analyses described in this manuscript were obtained from dbGaP at http://www.ncbi.nlm.nih.gov/gap through dbGaP accession number phs000424.v8.pht002741.v8.p2.

## Author Contributions

A.Q.F. designed the study. M.B.B., E.A.M. and A.Q.F. developed the tools. J.K., M.B.B., and A.Q.F. analyzed the data. J.W. and X.W. contributed to interpretation of data. A.Q.F. wrote the manuscript with input from the co-authors.

---

Supplementary Table 1. **The breakdown of trio types inferred by MRPC-LOND across GTEx tissues and cell types**.

Supplementary Table 2. **Number of principal components (PrCs) associated with trios across tissues**.

Supplementary Table 3. **The breakdown of trio types inferred by MRPC-ADDIS across GTEx tissues and cell types**.

Supplementary Table 4. **Summary of cis- and trans-gene mediation trios across GTEx tissues and cell types inferred by MRPC-LOND.**

Supplementary Table 5. **Cis-gene mediation trios with summary statistics across tissues inferred by MRPC-LOND**.

Supplementary Table 6. **Trans-gene mediation trios with summary statistics across tissues inferred by MRPC-LOND**.

Supplementary Table 7. **Cis-gene mediation trios with summary statistics across tissues inferred by MRPC-ADDIS**.

Supplementary Table 8. **Trans-gene mediation trios with summary statistics across tissues inferred by MRPC-ADDIS**.

Supplementary Table 9. **Summary of gene types of mediators**.

Supplementary Table 10. **HiC results in four tissues for trans-gene mediation trios inferred by MRPC-LOND**.

Supplementary Table 11. **HiC results in four tissues for trans-gene mediation trios inferred by MRPC-ADDIS**.

Supplementary Table 12. **Mediation trios inferred by the GMAC method**.

Supplementary Table 13. **Inference agreement between GMAC and MRPC-LOND on tested trios in five selected tissues**.

Supplementary Table 14. **Inference agreement between GMAC and MRPC-ADDIS on tested trios in five selected tissues**.

Supplementary Table 15. **Recall and precision of MRPC-ADDIS and MRPC-LOND on simulated data**.

Supplementary Table 16. **Recall and precision of MRPC and GMAC on simulated data**. Here, the two metrics are evaluated for the edge between the two genes (T_1_ and T_2_) in a trio. The five basic models are grouped into two classes: M_1_, M_2_ and M_4_ have an edge between the two genes, whereas M_0_ and M_3_ do not. We evaluated recall and precision for the two classes separately.

**Supplementary Figure 1.**
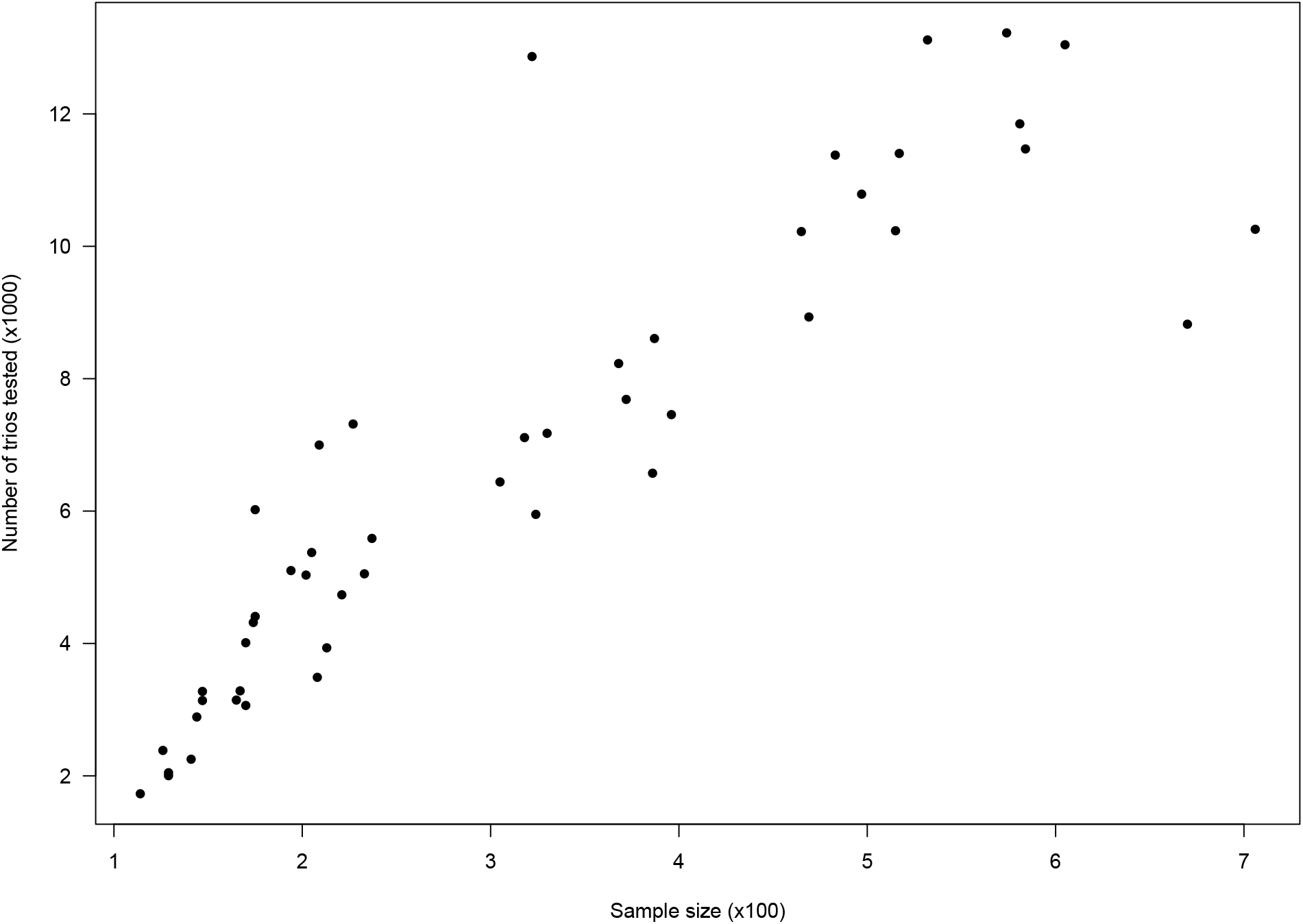
The number of trios tested versus the sample size in each tissue or cell type of the GTEx consortium.

**Supplementary Figure 2.**
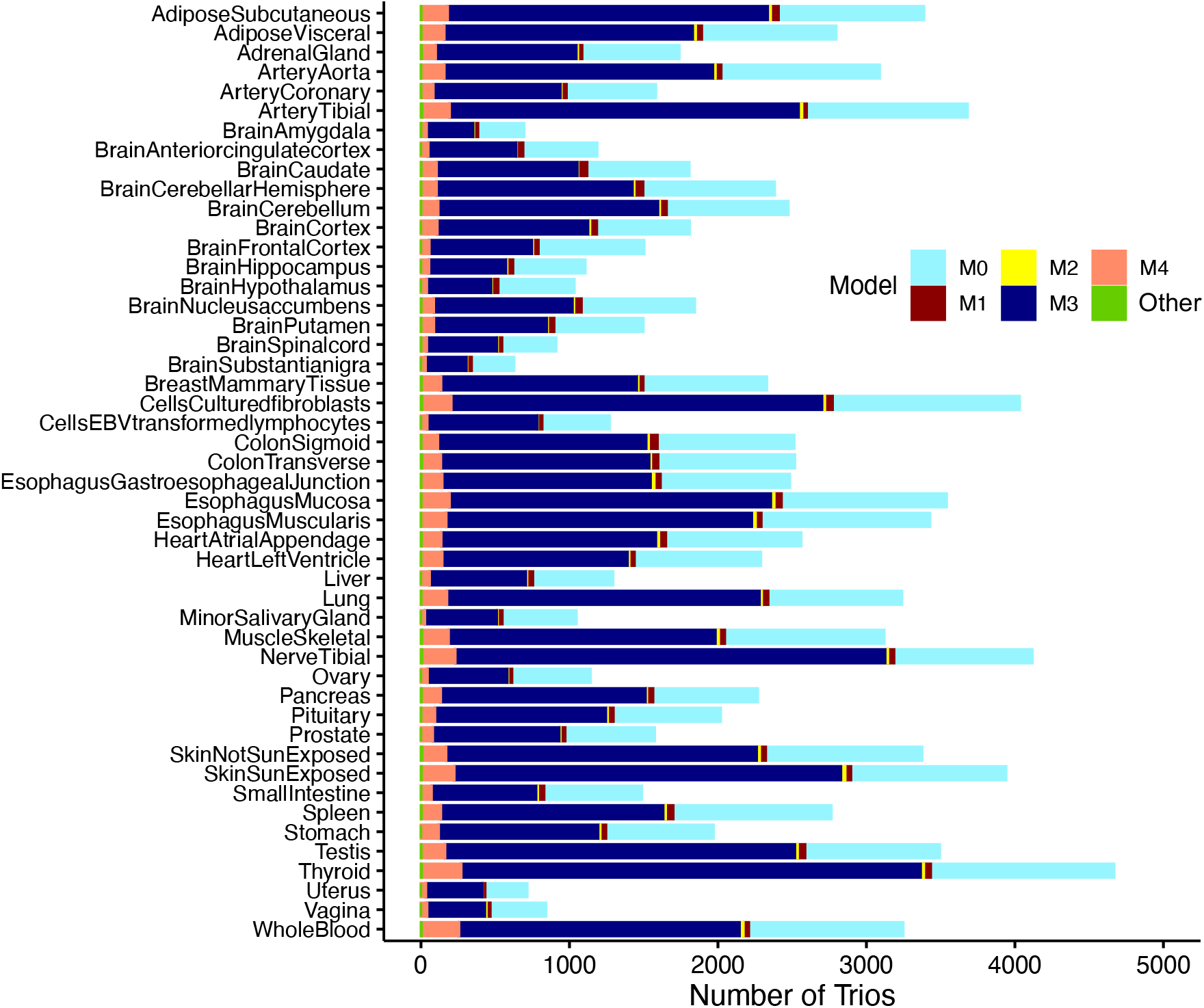
A stacked bar plot of MRPC-ADDIS inferred regulatory types of trios across GTEx tissues and cell types.

**Supplementary Figure 3.**
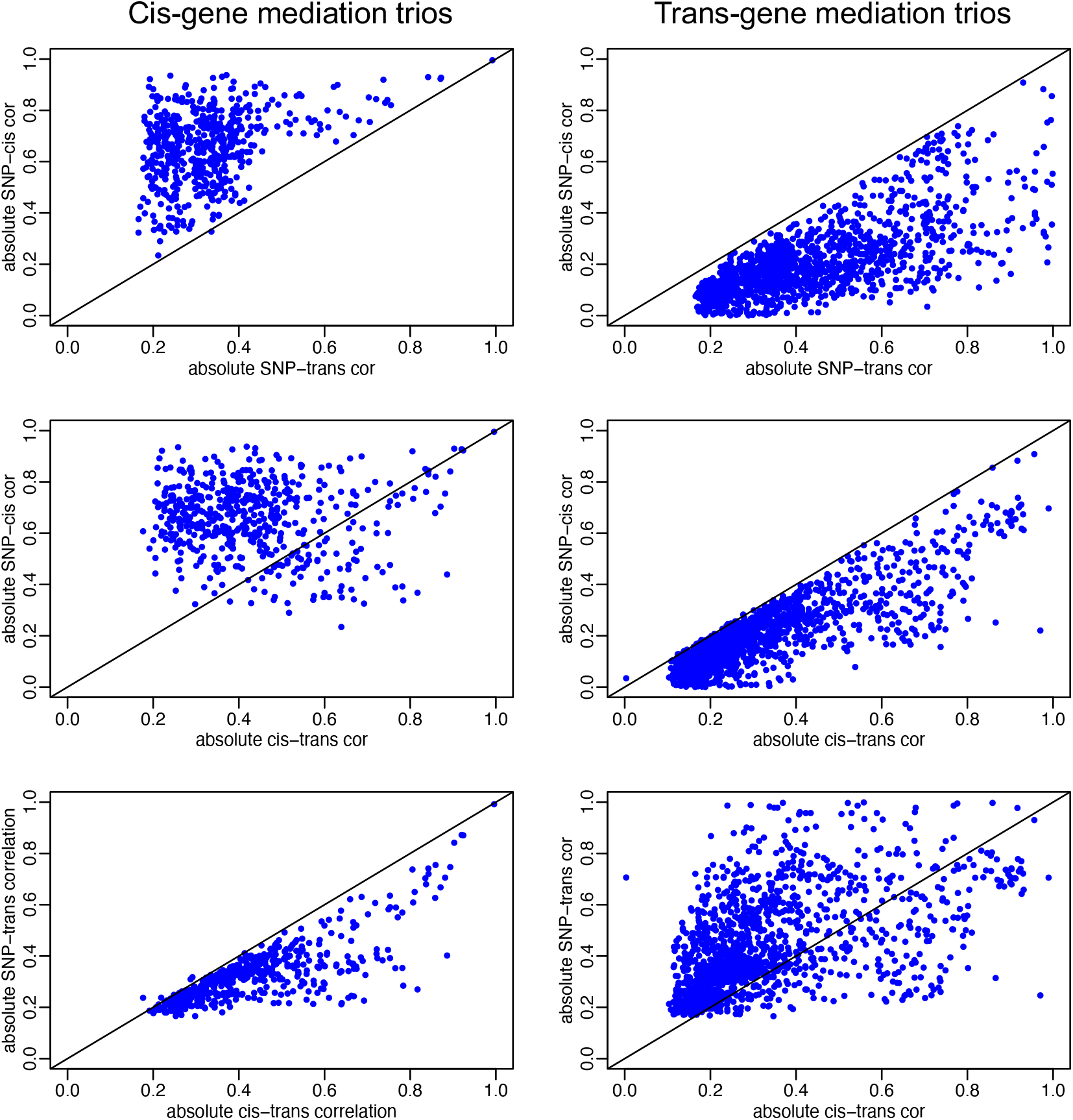
Correlations in MRPC-ADDIS inferred mediation trios.

**Supplementary Figure 4.**
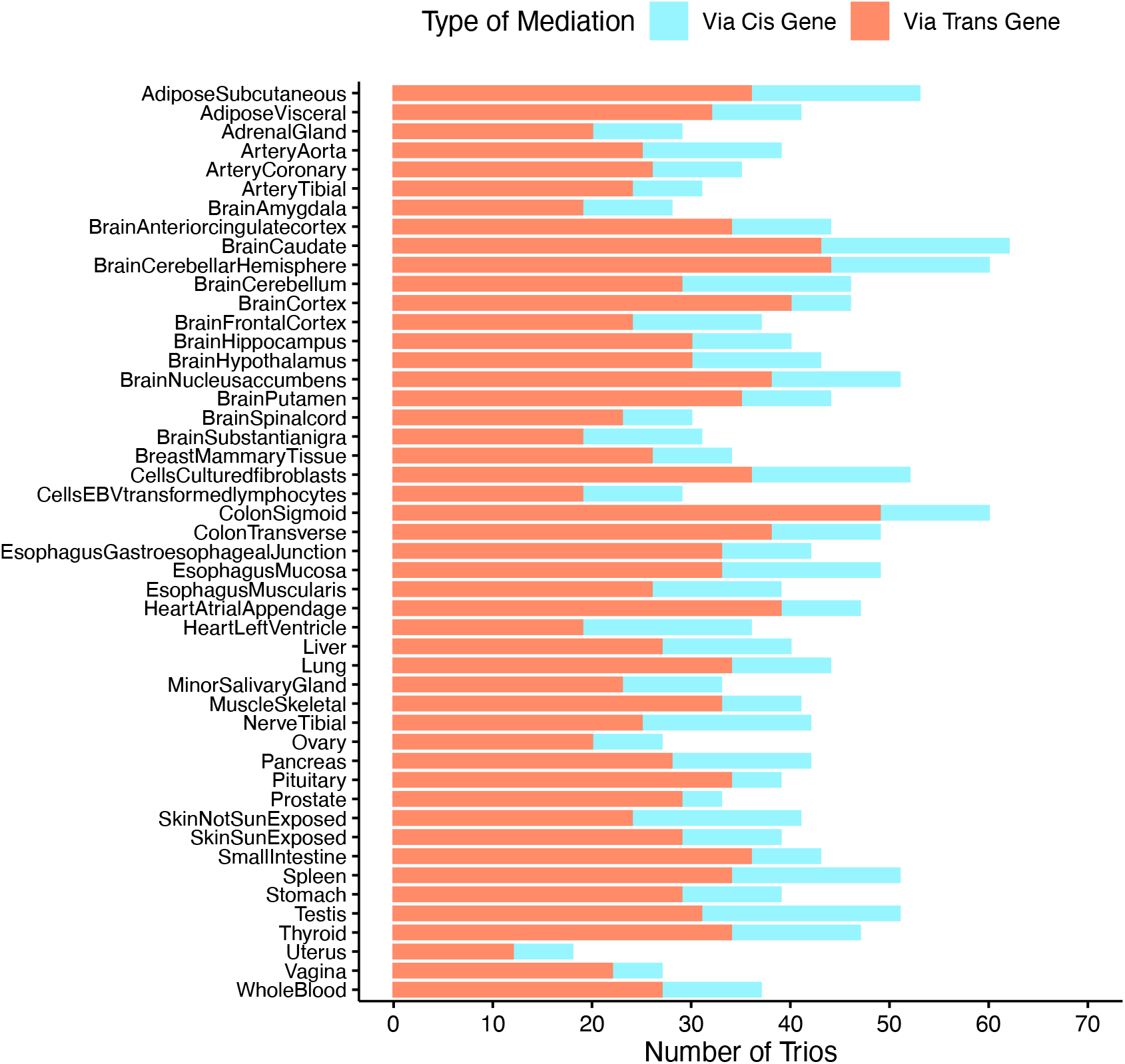
A stacked bar plot of MRPC-ADDIS inferred mediation types across GTEx tissues and cell types.

**Supplementary Figure 5.**
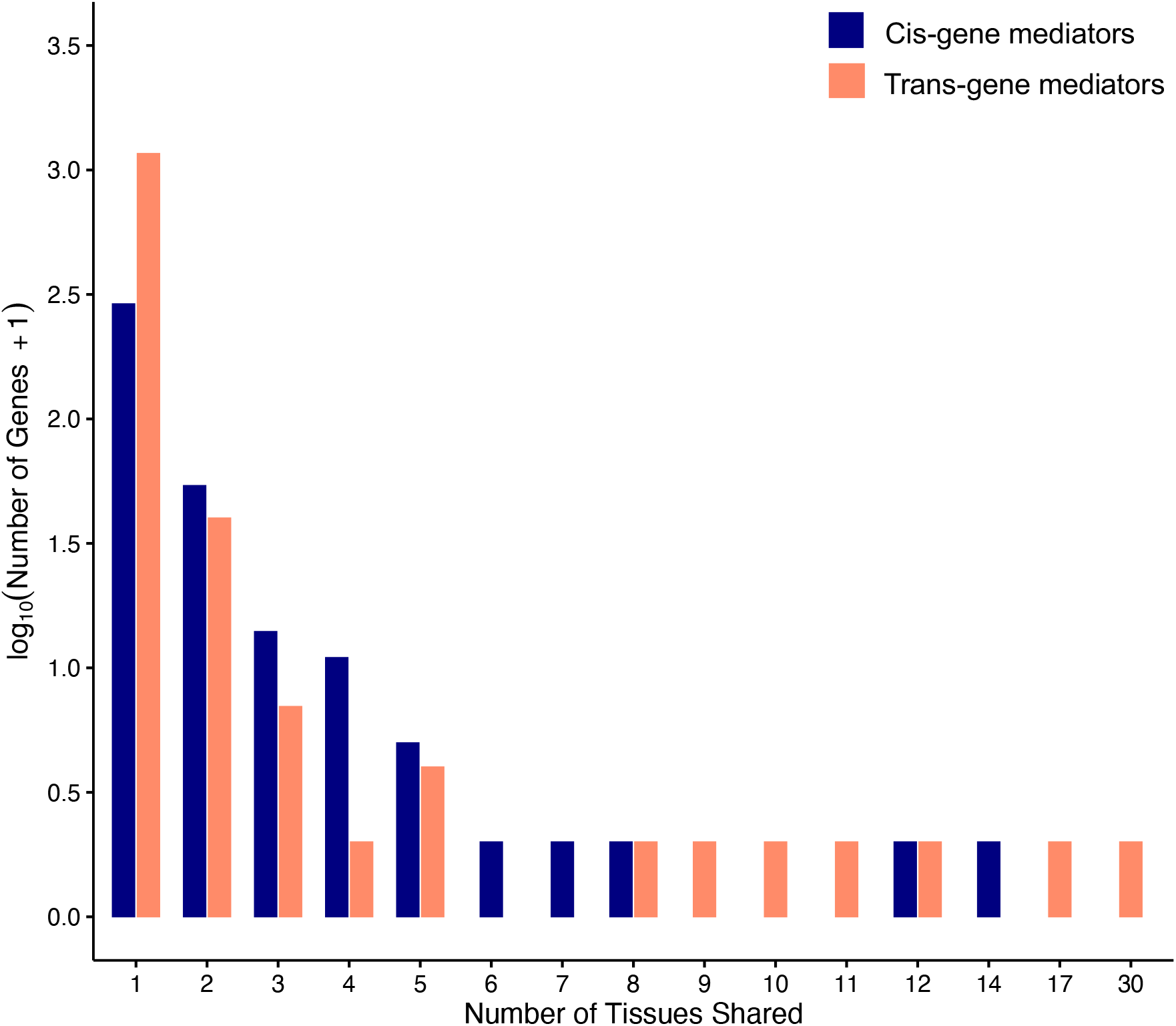
Histograms of tissue sharing for cis-gene and trans-gene mediators inferred by MRPC-ADDIS.

**Supplementary Figure 6.**
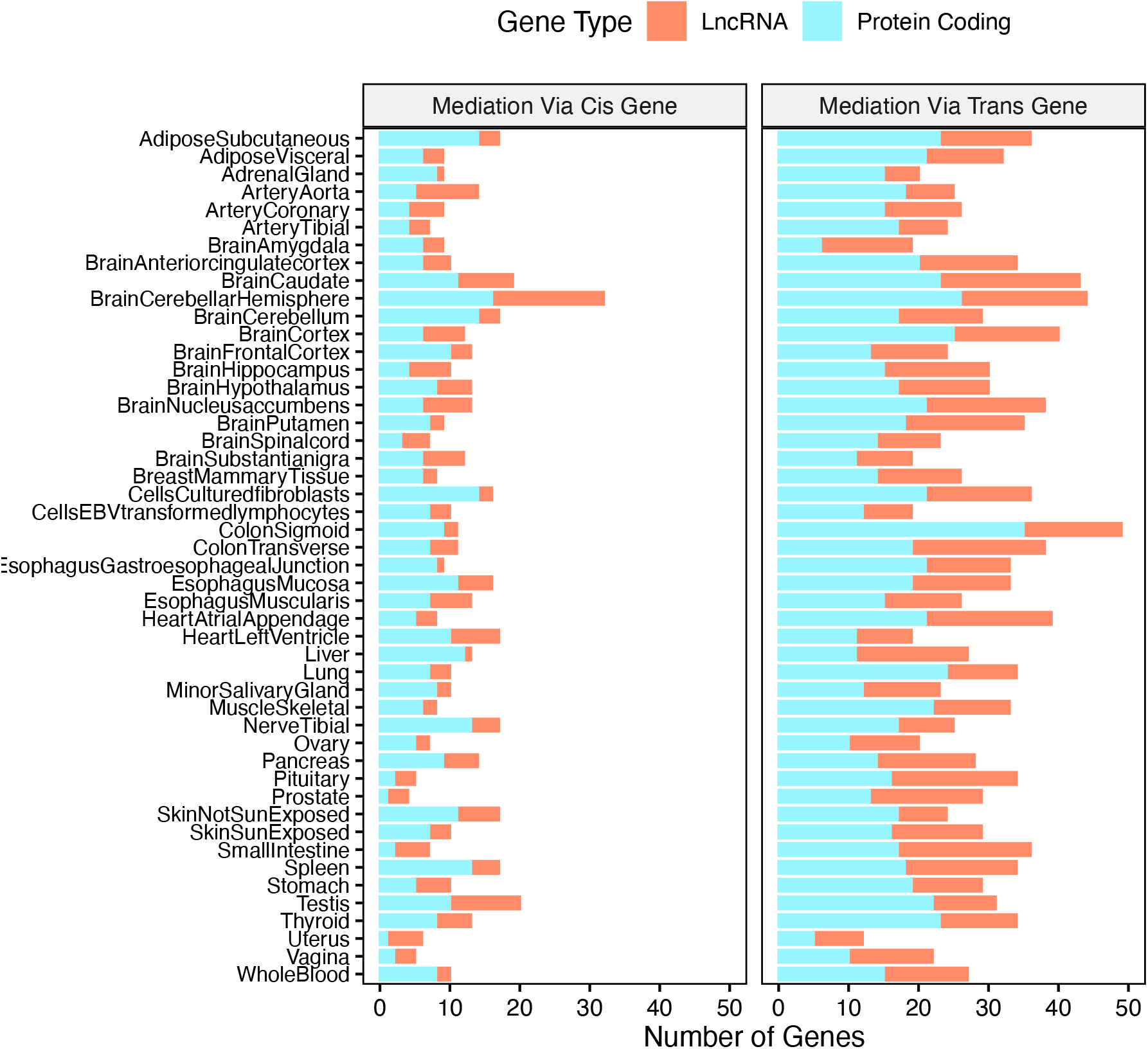
A stacked bar plot of MRPC-ADDIS inferred mediators by gene types.

## Notes

### Competing Interest Statement

The authors have declared no competing interest.

### Summary of Updates

The analysis is now focused on protein-coding genes and lncRNAs.

